# Studying vertical microbiome transmission from mothers to infants by strain-level metagenomic profiling

**DOI:** 10.1101/081828

**Authors:** Francesco Asnicar, Serena Manara, Moreno Zolfo, Duy Tin Truong, Matthias Scholz, Federica Armanini, Pamela Ferretti, Valentina Gorfer, Anna Pedrotti, Adrian Tett, Nicola Segata

**Affiliations:** Centre for Integrative Biology, University of Trento, Italy; Azienda Provinciale per i Servizi Sanitari, Trento, Italy

**Author notes:** Equal contribution (Francesco Asnicar, Serena Manara).

## Abstract

The gut microbiome starts to be shaped in the first days of life and continues to increase its diversity during the first months. Several investigations are assessing the link between the configuration of the infant gut microbiome and infant health, but a comprehensive strain-level assessment of vertically transmitted microbes from mother to infant is still missing. We longitudinally collected fecal and breast milk samples from multiple mother-infant pairs during the first year of life, and applied shotgun metagenomic sequencing followed by strain-level profiling. We observed several specific strains including those from *Bifidobacterium bifidum*, *Coprococcus comes*, and *Ruminococcus bromii*, that were present in samples from the same mother-infant pair, while being clearly distinct from those carried by other pairs, which is indicative of vertical transmission. We further applied metatranscriptomics to study the *in vivo* expression of vertically transmitted microbes, for example *Bacteroides vulgatus* and *Bifidobacterium* spp., thus suggesting that transmitted strains are functionally active in the two rather different environments of the adult and infant guts. By combining longitudinal microbiome sampling and newly developed computational tools for strain-level microbiome analysis, we showed that it is possible to track vertical transmission of members of the microbiome from mother to infants and characterize their transcriptional activity. Our work poses the basis for surveying at larger scale the sources of microbial diversity in the infants and starts associating these transmissions with the subsequent longer-term development of a healthy or dysbiotic microbiome.

**Importance:** Early infant exposure is important in the acquisition and ultimate development of a healthy infant microbiome. There is increasing support that the maternal microbial reservoir is a key route of microbial transmission, yet much is inferred from the observation of shared species in mother and infant. Common species, *per se*, does not necessarily equate vertical transmission as species exhibit considerable strain heterogeneity and it is therefore imperative to identify shared strains. We demonstrate here the potential of shotgun metagenomics and strain-level resolution to identify vertical transmission events via the maternal source. Combined with a metatranscriptomic approach, we show the potential not only to identify and track the fate of microbes in the early infant microbiome but also identify the metabolically active members. These approaches will ultimately provide important insights into the acquisition, development and community dynamics of the infant microbiome.

## Introduction

The community of microorganisms that dwell in the human gut has been shown to play an integral role in human health (1–3), for instance by harvesting nutrients that would be inaccessible otherwise (4), modulating host metabolism and immune system (5) and preventing infections by occupying the ecological niches that could otherwise be exploited by pathogens (6). The essential role of the intestinal microbiome is probably best exemplified by the successful treatment of dysbiotic states such as chronic life threatening *Clostridium difficile* infections using microbiome transplantation therapies (7–9).

The gut microbiome is a dynamic ecological community which is shaped by multiple factors throughout an individual’s life. The early development of the infant microbiome has been shown to be particularly crucial for future health (10–12) and a few studies have investigated the factors that are important in defining its first structure (13–16). In particular, gestational age at birth (16), mode of delivery (13, 14), early antibiotic treatments (17), have all been shown to influence the gut microbial composition in the short term, and the pace of its development in the longer term.

Vertical transmission of bacteria from the body and breast milk of the mother to her infant has gained attention as an important source of microbial colonization (13, 18–20). The extent to which the mother directly contributes to the infant’s gut microbial diversity is likely more crucial than microbial transmission from exposure to the wider environment (21, 22), including the delivery room (23). Early cultivation-based and cultivation-free methods (16S rRNA community profiling and a single metagenomic study) have suggested that the mother could transfer microbes to the infant by breastfeeding (24), and that the route of delivery has the potential of seeding the infant gut with members of the mother’s vaginal community (10, 13, 25, 26).

Studies of the vertical transmission of microbes from mothers to infants have however hitherto focused on the cultivable fraction of the community (27) or lacked strain-level resolution (10). At a species level resolution, where the same microbial species have been found to be shared between mothers and their infants (12, 28), it still remains inconclusive if vertical transmission has actually occurred as the same species are also observed in unrelated mothers and infants. Because different individuals will often carry different strains of common species (29, 30), it is crucial to profile bacteria at a finer strain-level to ascertain the route of transfer. This has been performed only for specific microbes by cultivation methods (15, 27), but many vertically transmitted microorganisms remain hard to cultivate (15), thus the true extent of microbial transmission remains unknown. A further crucial aspect still largely unexplored is the fate of vertically acquired strains i.e. are they merely transient or functionally active and therefore more likely to be early colonizers of the infant intestine. Although studies have reported the transcriptional activity of intestinal microbes under different conditions (31-34), no studies have applied metatranscriptomics to characterize the activity of vertically transmitted microbes *in vivo*.

In this work, we present and validate a shotgun metagenomic pipeline to track mother-to-infant vertical transmission of microbes by applying the strain-level computational profiling to all the members of the infant microbiome. Moreover, we assessed the transcriptional activity of vertically transmitted microbes to elucidate if transferred strains are not only present but also metabolically active in the infant gut.

## Results and Discussion

We analyzed the vertical transmission of microbes from mother-to-infant by enrolling 5 mother-infant pairs and collecting fecal samples and breast milk (see **Methods**) at three months of age of the infant (Timepoint 1). Two mother-infant pairs were additionally sampled at 10 months (Timepoint 2) and one pair at 16 months post birth (Timepoint 3). We applied shotgun metagenomic sequencing to 24 microbiome samples (8 mother fecal samples, 8 infant fecal samples, and 8 milk samples) generating 1.2G reads (39.6M s.d. 28.7M reads/sample, see **Supplementary Table S1**). Metatranscriptomics (90.55M s.d. 46.86M reads/sample) was also applied on fecal samples of two pairs to investigate the differential expression profiles of the bacterial strains in the gut of mothers and their infants.

### Shared mother-infant microbial species

In our cohort, the infant intestinal microbiome is dominated by *Escherichia coli* and *Bifidobacterium* spp., such as *B. longum*, *B. breve* and *B. bifidum* (Fig. 1A), which is consistent with what has been described previously (11, 35, 36). As expected, mothers’ intestines have a more diverse microbiome with high abundances of *Prevotella copri*, *Clostridiales* (e.g. *Coprococcus* spp. and *Faecalibacterium prausnitzii*), and *Bacteroidales* (e.g. *Parabacteroides merdae* and *Alistipes putredinis*). Interestingly, the post-weaning infant microbiome of Pair 5 (Timepoint 3, 16 months post birth), had already shifted towards a more “mother-like” composition (Fig 1B), with an increase in diversity and the appearance of *Parabacteroides merdae*, *Coprococcus* spp., and *Faecalibacterium prausnitzii* (12, 36). Nevertheless, this 16-months old infant still retains some infant microbiome signatures, such as a high abundance of *Bifidobacteria* that are present only at low levels in mothers’ samples (Fig. 1A and 1C).

**Figure 1.**
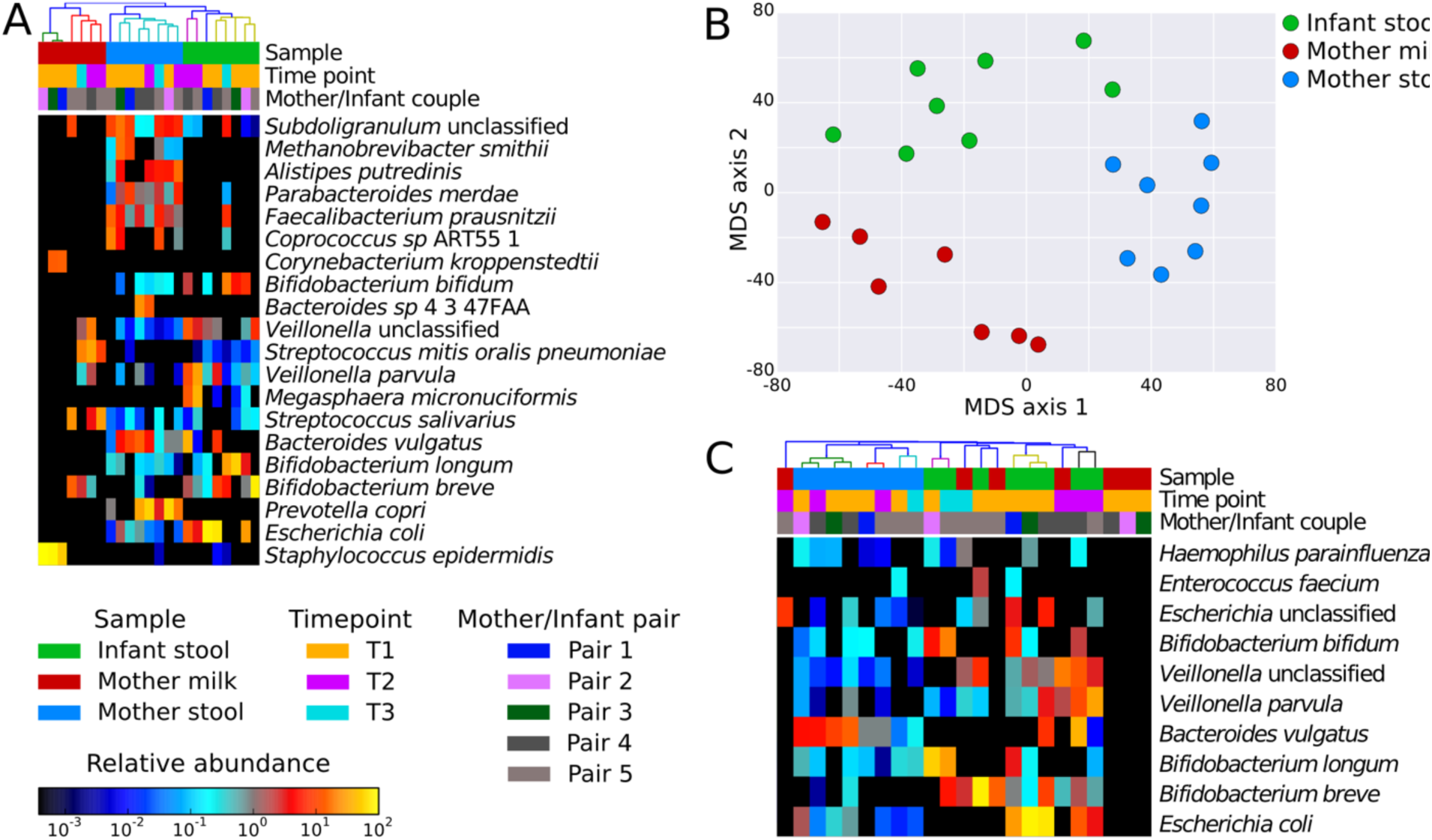
Microbial composition of mother and infant samples, and shared bacteria within mother-infant pairs. **(A)**Quantitative microbial taxonomic composition of the metagenomic samples from milk and fecal samples of mothers and infants as estimated by MetaPhlAn2 analysis (45) (only the 20 most abundant species are indicated). As can be seen milk samples present a low microbial richness compared to fecal samples. **(B)** Ordination plot of microbiome composition showing clustering of the three different sample types: mother feces, infant feces, and breast milk samples. The two infant samples close to the cluster of mother feces and in between the clusters of mothers and infants, are from later timepoints, denoting the convergence of the infant microbiome towards an adult-like one. **(C)** The abundances of the 10 microbial species detected (>0.1% abundance) in at least one infant and their respective mother (shared species have been identified based on samples from Timepoint 1 only).

We extracted and successfully sequenced microbial DNA from 7 out of 8 milk samples. Profiling of the microbial composition of milk is hindered by the high abundance of interfering molecules (proteins, fats, proteases -e.g. plasmin and calcium ions) (37–39) that affect the efficiency of extraction and amplification steps. Even so, we obtained an average of 3.08 s.d. 1.5 Gb (100 bp reads) per sample. In each sample we extracted 26 s.d. 56 Mb of non-human reads (higher than the only other metagenomic study (40), see **Supplementary Table S1**).

Milk samples had a limited microbial diversity at the first sampling time (Timepoint 1, three months post birth), and included skin-associated bacteria like *Corynebacterium kroppenstedtii* and *Staphylococcus epidermidis*. Cutaneous taxa, however, were only observed in low abundancies in the gut microbiome of infants, confirming that skin microbes are not colonizers of the human gut (Fig. 1A). At later timepoints the milk samples were enriched in *B. breve* and bacteria usually found in the oral cavity, such as *Streptococcus* and *Veillonella* spp. The presence of oral taxa in milk has already been observed by previous studies based on 16S rRNA sequencing (13, 19, 20, 24) and shotgun metagenomics (40). This could be caused by the retrograde flux into the mammary gland during breastfeeding (41) whereby cutaneous microbes of the breast and from the infant oral cavity are transmitted to the breast glands (42). These observations are summarized in the ordination analysis (Fig. 1B), in which the different microbiome samples (infant feces, mother feces, and milk) clustered by type, with weaning representing a key factor in the shift from an infant to an adult-like microbiome structure (12, 36, 43).

Comparing the species present both in the mother and infant pairs (Fig. 1C), we observed that many shared species occur at much higher abundance in the infant than in the mother (e.g. *Escherichia*, *Bifidobacterium* and *Veillonella* spp.) possibly due to the lower level of species diversity and therefore competition in the gut. *Bacteroides vulgatus* was instead found at relatively high abundances in both the infant and the mother of Pair 4 at both Timepoint 1 and Timepoint 2. The presence of shared species in infant-mother pairs seen here and elsewhere (13, 15, 16, 24, 44) confirms that mothers are a potential reservoir of microbes vertically transmissible to infants, but it remains unproven whether the same strain is transmitted to the infant from the mother or if a more complex transmission route is involved.

### Strains shared between mother and infants are indicative of vertical transmission

At a species level resolution, it has been shown that different individuals have a core of microbial species, but these common species consist of distinct strains (29, 30). To analyze microbial transmission from mother to infant it is therefore crucial to assess whether they carry the same microbial strain for that species. To this end, we further analyzed the metagenomic samples at a finer strain level resolution. This was achieved by applying a recent strain-specific pangenome-based method called PanPhlAn (29) as well as a novel method, called StrainPhlAn (see **Methods**), that identifies single-nucleotide variants (SNVs) in species-specific marker genes.

Using this SNVs-based analysis, we first confirmed considerable strain-level heterogeneity in the species present in multiple mothers also with respect to available reference genomes (Fig. 2 and **Fig. S2**). This heterogeneity is not observed between mother-infant pairing, as in the case of *Bifidobacterium* spp., *Ruminococcus bromii*, and *Coprococcus comes*. The infant of Pair 4 at Timepoint 2 harbors, for example, a strain of *B. bifidum* which matches the one in his mother at 99.96% sequence identity and yet at the same time is clearly distinct from the *B. bifidum* strains harbored by other infants that differ by at least 0.64% nucleotides (the average nucleotide diversity with the other strains in other pairs is 0.74%, Fig. 2A). The same is true for the strains of *C. comes* (99.87% intra-pair similarity, 0.87% and 1.63% divergence compared to the closest strain and average, respectively, Fig. 2B) and *R. bromii* (99.93% similar and 1.53% and 2.63% diverse – same as above, Fig. 2C) that are shared by Pair 5. The mother-infant sharing of the same strain was confirmed by strain-level pangenome analysis (29) that showed that the strains from the same pair carry the same unique gene repertoire (**Supplementary Fig. S3**). While it is not possible to exclude the independent acquisition of strains from a shared environmental source, the finding that mother-infant pairs harbor some shared strains represents strong evidence of vertical microbiome transmission.

**Figure 2.**
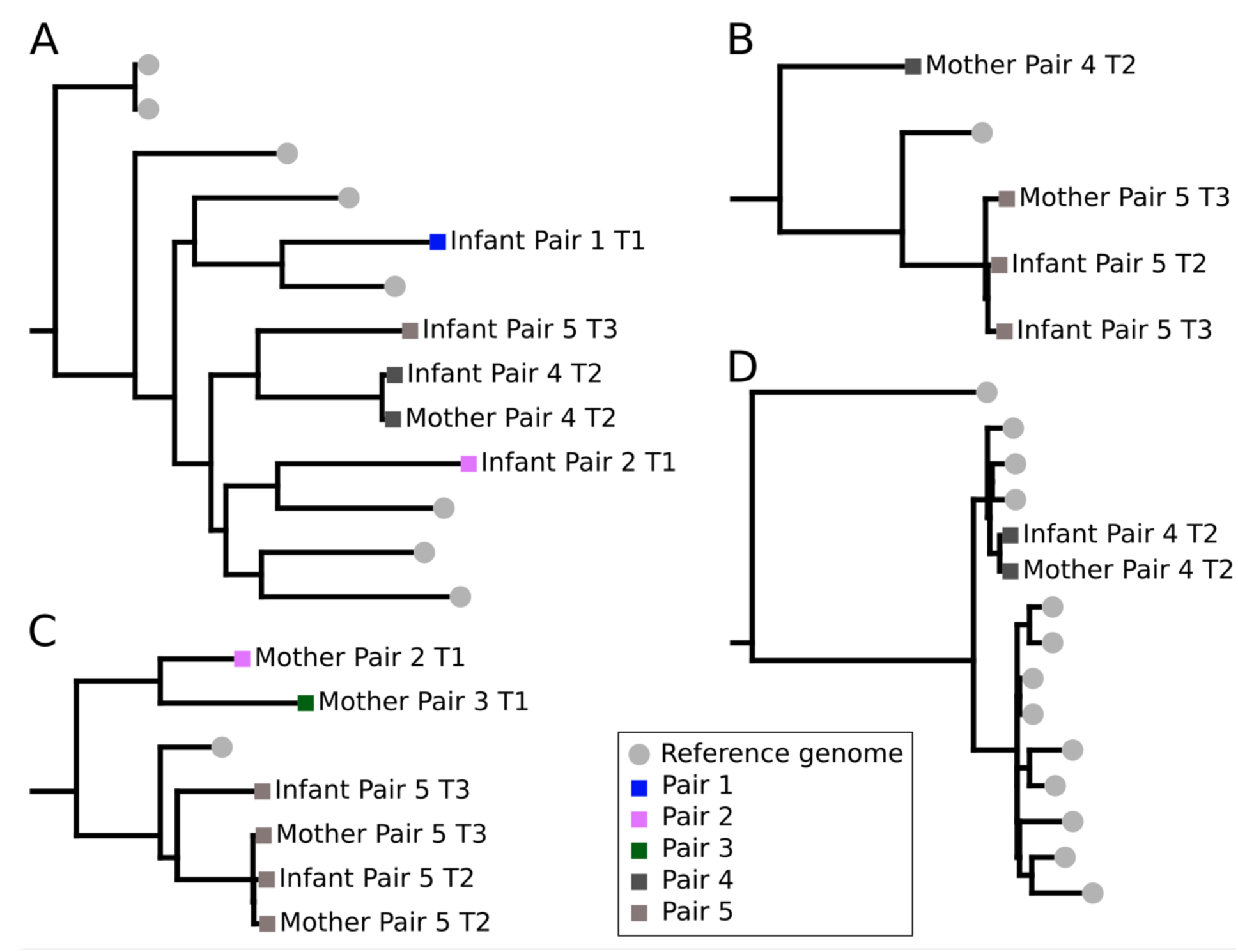
Strain-level phylogenetic trees for microbes present in both the mother and infant. Phylogenetic trees built by using species-specific markers (StrainPhlAn, see **Methods**) confirming the presence of the same strain in the mother and infant intestinal microbiome, thus suggesting vertical transmission. Available reference genomes were included in the phylogenetic trees. Here we report three bacterial species, namely **(A)***Bifidobacterium bifidum*, **(B)***Coprococcus comes*, and **(C)***Ruminococcus bromii*, and the most abundant viral species found in Pair 4 **(D)** Pepper mild mottle virus. Other species-specific phylogenetic trees (*B. adolescentis*, *B. breve*, and *B. longum*) are reported in the **Supplementary Figure S2**.

The mother-infant Pair 5 at Timepoint 3 harbors the highest number of shared species with 35.2% of the species present in the infant microbiome at a relative abundance >0.1% that are also present in the mother according to MetaPhlAn2 profiles. Among the 35.2% of shared species, the mother and infant of Pair 5 shared several strains (Fig. 2 and **Supplementary Fig. S2**), according to the results of both PanPhlAn and StrainPhlAn (see for example *B. adolescentis* and *C. comes*, **Fig. 2** and **Supplementary Fig. S2** and **S3**). However, some strains that were shared within mother-infant Pair 5 at earlier timepoints were replaced by different ones at Timepoint 3. The *R. bromii* strain harbored by the infant at Timepoint 3 is in fact different from the one found at Timepoint 2 and in the mother at both timepoints (Fig. 2C) and a similar event is observed for the latter infant timepoint for *B. breve* (**Supplementary Fig. S2B**) and *B. longum* (**Supplementary Fig. S2C**). Although it is not possible to generalize these results because of the small sample size, these replacement events suggest that the originally acquired maternal strains can later be substituted with different ones from the same species (46, 47).

We then extended our analysis to the viral organisms detectable from metagenomes and metatranscriptomes, as viruses also can be potentially vertical transmitted. The DNA viruses identified from our metagenome samples largely consisted of bacteriophages from the *Caudovirales* order, a common order found in the intestine as previously reported (3, 48), but the low breadth of coverage for many of them made it difficult to identify pair-specific phage variants (**Supplementary Table ST3**). When looking at RNA viruses from metatranscriptomics, we detected the Pepper Mild Mottle Virus (PMMoV), a single-stranded positive-sense RNA virus of the genus *Tobamovirus*, in all of the four metatranscriptomes from pairs 4 and 5. Surprisingly, transcripts from the PMMoV where in greater abundance than all the other microbial transcripts found for the mother of Pair 4. PMMoV has already been reported in the gut microbiome (49-51), and other viruses of the same family have been shown to be able to enter and persist in eukaryotic cells (52, 53). The high abundance of PMMoV in mother-infant Pair 4 allowed us to reconstruct its full genome (99.9%) and to perform a phylogenetic analysis showing that both the mother and the infant share a PMMoV with the very same genomic sequence, which is highly similar but clearly distinct from the PMMoV reference genomes (27 SNVs in total, Fig. 2D). Although the coverage was lower, the same evidence of a shared PMMoV was observed also within Pair 5. The analysis of PMMoV polymorphisms within each sample also suggests the coexistence of different PMMoV haplotypes (**Supplementary Figure S6**) in the same host. Although vertical transmission of RNA viruses and PMMoV specifically would be intriguing, because of the age and dietary habits of the infants (**Supplementary Table ST1**) this finding could be related to the exposure to a common food source (54).

### Differences in the overall functional potential and expression in mothers and infant

The physiology of the mammary gland (milk) as well as the adult and infant intestine is reflected by niche specific microbial communities. We complemented the taxonomic analysis above by employing HUMAnN2 (see **Methods**) to characterize the overall functional potential of these inhabiting communities. As expected, there was considerably overlap in the functionality of the mothers’ and infants’ gut microbiomes when looking at the relative abundance of microbial pathways (Fig. 3A). Nevertheless, there were notable differences, for instance the infants’ microbiome showed a higher potential for utilization of intestinal mucin as a carbon source and for folate biosynthesis, (11, 55, 56) while displaying a lower potential for starch degradation, in line with what had been observed previously (11, 55–58). Mucin utilization specifically by infant-gut microbial communities is reflective of the higher abundance of mucin degrading *Bifidobacteria* observed from the taxonomic analyses (11, 55–57), whereas increased folate biosynthesis (11, 55, 56, 58) and decreased starch degradation (4) have been suggested to be a response to the limited dietary intake in infants with respect to adults. Interestingly, the post-weaning infant of Pair 5 (16 months post birth) clustered together with the adults’ intestinal samples (Fig. 3B), suggesting that the shift towards an adult-like microbiome observed in the taxonomic profiling (Fig. 1B) is also reflected by or is as a consequence of a change in community functioning. Among the most prevalent pathways in the milk microbiomes we observed those involved in galactose and lactose degradation (59) as well as in aromatic compounds biosynthesis (**Supplementary Fig. S4**). This is specifically true for chorismate production, a key intermediate for the biosynthesis of essential amino acids and vitamins found in milk (58) (Fig. 3C and **Supplementary Fig. S4**).

**Figure 3.**
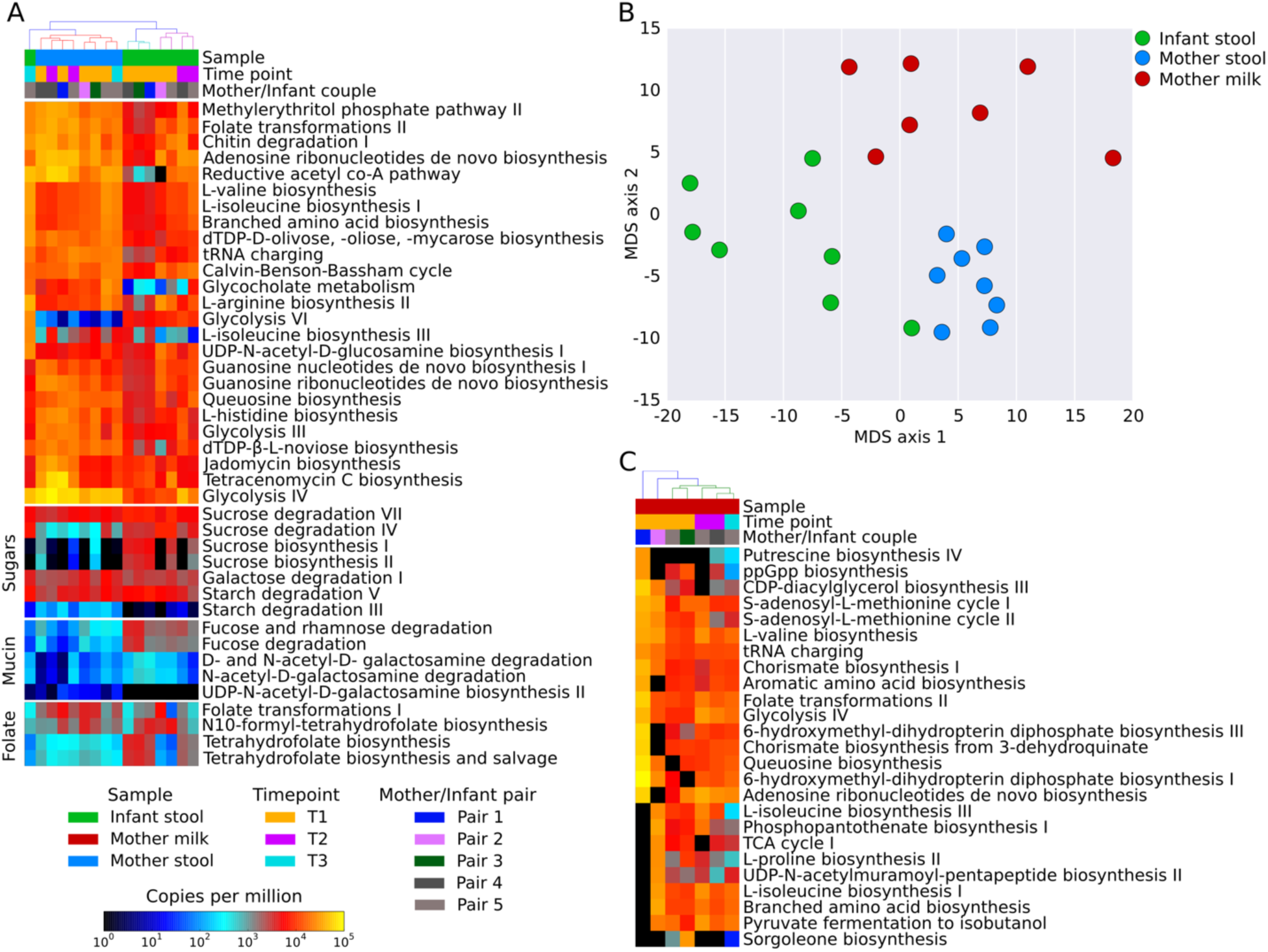
Functional potential analyses. **(A)** HUMAnN2 heatmap reporting the 25 most abundant pathways in the fecal samples of mothers and infants. Added at the bottom, specific pathways of interest: starch, mucin, and folate metabolism. **(B)** MDS result on functional potential, showing the differences between fecal samples of mothers and infants, and milk. In particular, the infant-feces point in the mother-feces cluster corresponds to the Timepoint 3 of the Pair 5, showing a shift from the infant microbiome towards and adult-like microbiome. **(C)** HUMAnN2 results on the 25 most abundant pathways found only in the milk samples.

To further evaluate the functional capacity of the gut associated microbiomes and analyze the *in vivo* transcription, we performed metatranscriptomics on the feces of two mother-infant pairs (see **Methods**). HUMAnN2 was used to identify differences in the transcriptional levels of pathways in mothers and infants gut. The most notable global difference was that fermentation pathways were highly transcribed in the mother metatranscriptome compared to infant. This reflects the transition of the gut from aerobic to anaerobic and the associated shift from facultative anaerobes to obligate anaerobes over the first few months of life (60, 61). The same is true for pathways involved in starch degradation, which are not only poorly represented but also negligibly expressed in the infants’ microbiomes (59). What is evident is that the transcriptional patterns for different members vary considerably, as illustrated for Pair 4 and 5 (Fig. 4A, and **Supplementary Fig. S5A** respectively). For example, in Pair 4 we observe in the infant that *B. vulgatus* is more transcriptionally active (average of 2.7 s.d. 2.5 normalized transcript abundance – NTA, see **Methods**) than both *E. coli* (average 0.4 s.d. 0.6 NTA) and *Bifidobacteria* spp. (average 0.01 s.d. 0.01 NTA), although whether these differences are physiologically significant remains less clear.

**Figure 4.**
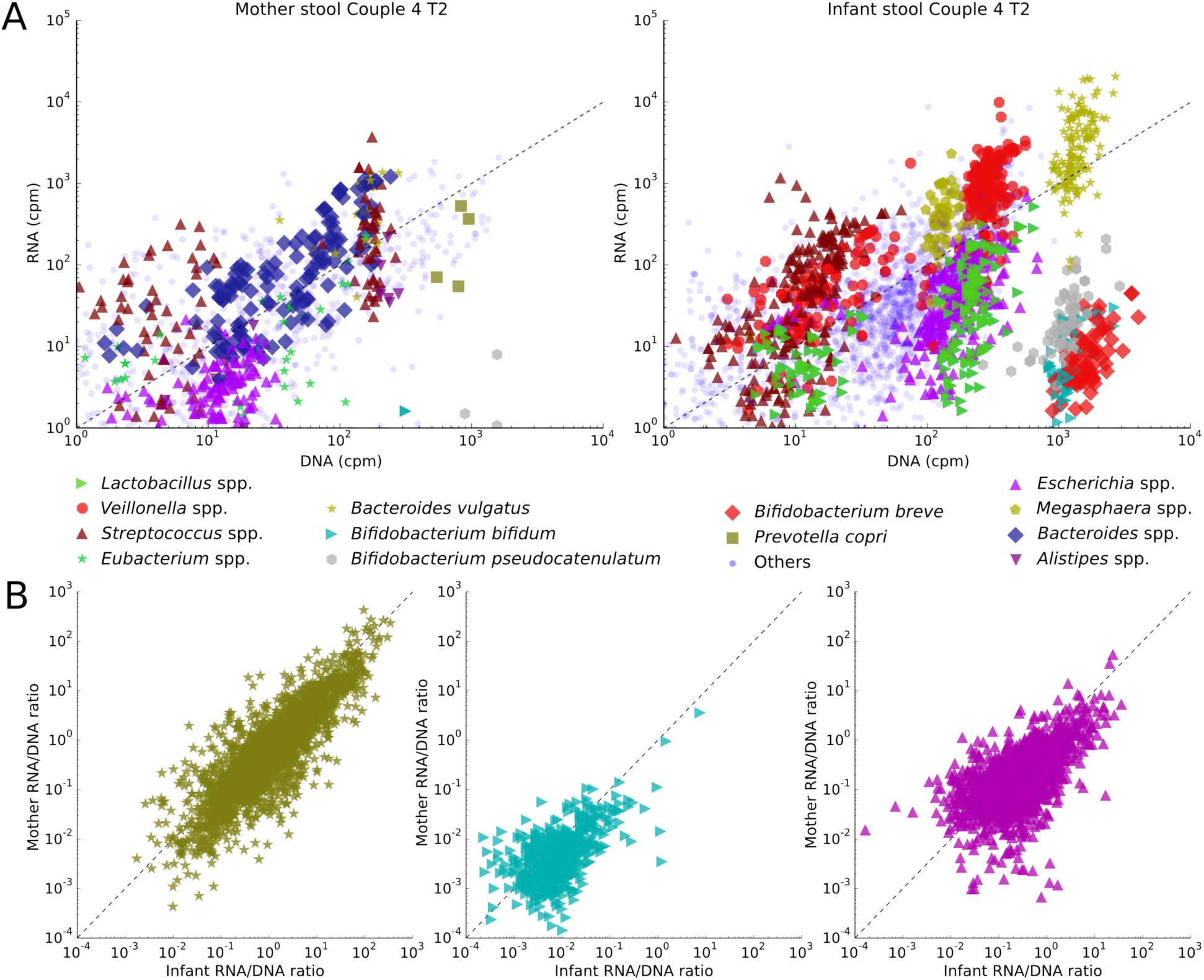
Metabolic pathways expression in mother and infant of Pair 4 at Timepoint 2. Scatterplots showing the transcription rate of metabolic pathways of different species and genera of interest for both the mother and infant of Pair 4 at Timepoint 2. (B) Comparison between gene families’ transcription rate in mother and infant gut microbiomes.

### Strain-specific transcriptional differences in mothers and infants

To further explore the transcriptional activity of the intestinal microbiomes and more specifically to ascertain which individual microbial members are metabolically active in the gut, we employed the strain-specific metatranscriptomic approach implemented in PanPhlAn (29) (see **Methods**). Of particular interest is the transcriptional activity of the shared mother-infant strains that based on our strain-level analyses are likely to have been vertically acquired by the infant from a maternal route. Such transcriptional analyses can clarify whether these transmitted strains are not only present but also functioning in the infant gut suggesting that the transmitted strains are likely colonizing. For three transmitted species in Pair 4 (*B. vulgatus, E. coli* and *B. bifidum*) we show that they are both active in the mother and infant intestine (Fig. 4B). Of note is *B. bifidum* that is more active in the infant with respect to the mother (Fig. 4B), which would be expected as this species is a known early colonizer of the infant gut (11, 35, 36). Interestingly, the *B. bifidum* strain of Pair 5 shows the opposite behavior (Supplementary Fig. S5B). We postulate that this is because the infant of Pair 5 was post weaning (10% breast milk diet) compared to the pair 4 (90% breast milk diet) and reflects the change in substrate availability from breast milk to solid food, which might have a detrimental effect on the bifidobacterial population (36, 62, 63). Moreover, in support of our metagenomics analyses that the microbiome of infant of Pair 5 is shifting towards a more adult like structure (Fig. 1B), we observe high transcriptional activity for *R. bromii,* a commonly adult associated species, which could be seen as hallmark of this transition (64, 65).

It is well established that metatranscriptomic profiling provides a more accurate account of the actual community functioning than metagenomics alone. Here we show that the combination of both approaches affords the exploration of which members are not only transmitted but are also actively participating in the community, and therefore a more detailed account of the microbial community dynamics.

## Conclusions

Human associated microbiomes are complex and dynamic communities that are continuously interacting with each other, their host, as well as under influence from environmental sources of microbial diversity. Identifying and understanding the transmission from these external sources is crucial to understand, in particular, how the infant gut microbiome is colonized and ultimately develops into an adult like composition. However, detecting direct transmission is not a trivial task: many species are ubiquitous in host-associated and in the wider environment alike, yet they actually comprise a myriad of different strains and phenotypic capabilities. Therefore, detection of microbial transmission events requires the ability to characterize microbes at the strain level. The epidemiological tracking of pathogens from cultivation-based isolate sequencing has proven successful (66, 67) but it relies on time-consuming protocols and can focus only on a limited number of species. In contrast, while there have been some examples of strain level tracking from metagenomic data (67, 68), this still remains challenging. In this study, we developed methods for identifying the vertical flow of microorganisms from mothers to their infants and show that maternal microbial sources are important in the development of the infant gut microbiome.

We demonstrate that high-resolution computational methods applied to shotgun metagenomic and metatranscriptomic data enable the tracking of strains and the following of strain-specific transcriptional patterns across mother-infant pairs. In our cohort of five mother-infant pairs, we detected several species with substantial genetic diversity between different pairs but identical genetic profiles in the mother and her infant, indicative of vertical transmission. These include some *Bifidobacteria* typical of the infant gut (i.e. *B. longum*, *B. breve*, *B. bifidum*, and *B. adolescentis*) but also *Clostridiales* usually found in the adult intestine (i.e. *R. bromii* and *C. comes*) and viral organisms. These results thus confirm that the infant receives a maternal microbial imprinting that is likely crucial for the development of the gut microbiome in the first years of life.

The strain-level investigation of vertical transmitted microbes was followed by transcriptional activity characterization of the transmitted strains in the mother and infant environments. We found that the transcriptional activity of shared strains within mother-infant pairs proved to be different in the two hosts, suggesting that successful colonization of maternally transmitted microbes.

Altogether, our work provides the preliminary results and methodology to expand our knowledge on how microbial strains are transmitted across microbiomes. Expanding the cohort size and considering other potential microbial sources of transmission, such as additional mother and infant body sites, as well as other family members (i.e. fathers and siblings) and environment (hospital and house surfaces), will likely shed light on the key determinants in early infant exposure and the seeding and development of the infant gut microbiome.

## Materials and methods

### Sample collection and storage

In total five mother-infant pairs were enrolled. Fecal samples and breast milk were collected for all pairs at three months (Timepoint 1), additional samples were collected for Pair 4 and 5 at 10 months (Timepoint 2) and for Pair 5 only at 16 months (Timepoint 3) (**Supplementary Table SX**). All aspects of recruitment, sample and data processing was approved by the local ethics committee. Fecal samples were collected in sterile Faeces tubes (Sarstedt, Nümbrecht, Germany) and immediately stored at -20°C. In those cases, where metatranscriptomics was applied, a fecal aliquot was removed prior to freezing the remaining feces. This aliquot was stored at 4°C and RNA extracted within two hours of sampling to preserve RNA integrity. Milk was expressed and collected mid flow by mothers into 15 ml centrifuge tubes (VWR, Milan, Italy) and immediately stored at -20°C. Within 48 hours of collection all milk/fecal samples were moved -80°C storage until processed.

### Nucleic acids extraction for metagenomic analysis

DNA was extracted from feces using the QIAamp DNA Stool Mini Kit (Qiagen, Netherlands), Milk DNA was extracted using the PowerFood Microbial DNA Isolation Kit (MO BIO, California, USA), both according to manufacturer’s specifications. Extracted DNA was purified using Agencourt AMPure XP kit (Beckman Coulter, California, USA). Metagenomic libraries where constructed using the Nextera XT DNA Library Preparation Kit (Illumina, California, USA) according to manufacturer instructions, and sequenced on the HiSeq 2500 platform (Illumina, California, USA) at an expected sequencing depth of 6 Gb/library.

### Nucleic acids extraction for metatranscriptomic analysis

Fecal samples for metatranscriptomic profiling were pretreated as described previously (69). Briefly, 110 μl of lysis buffer (30 mM Tris•Cl, 1 mM EDTA, pH 8.0, 1.5 mg/ml of proteinase K and 15 mg/ml of lysozyme) was added to 100 mg of feces and incubated at room temperature for 10 minutes. After pretreatment, samples were treated with 1200 μl of QIAGEN RLT Plus buffer (from AllPrep DNA/RNA Mini Kit, QIAGEN, Netherlands) containing 1% volume beta-mercaptoethanol, and transferred into 2 ml sterile screw-cap tubes (Starstedt, Germany) filled with 1 ml of zirconia-silica beads <0.1 mm (BioSpec Products, Oklahoma, USA). Tubes were placed on a Vortex-Genie 2 with Vortex Adapter 13000-V1-24 (MO BIO, California, USA) and shaken at maximum speed for 15 minutes. Lysed fecal samples were homogenized using QIAshredder spin columns (QIAGEN, Netherlands) and homogenized sample lysates were then extracted with the AllPrep DNA/RNA Mini Kit (QIAGEN, Netherlands) according to manufacturer’s specifications. Extracted RNA and DNA were purified using Agencourt RNAClean XP and Agencourt AMPure XP (Beckman Coulter, California, USA) kits respectively. Total RNA samples were subjected to rRNA depletion and metatranscriptomic libraries were prepared using the ScriptSeq Complete Gold Kit (Epidemiology) – Low Input (Illumina, California, USA). Metagenomic libraries were prepared with Nextera XT DNA Library Preparation Kit (Illumina, California, USA). All libraries were sequenced on the HiSeq 2500 platform (Illumina, California, USA) at an expected depth of 6 Gb/library.

### Sequencing data pre-processing

The metagenomes and metatranscriptomes were pre-processed by removing low quality reads (mean quality less than 25), trimming low quality positions (quality less than 15), and removing reads shorter than 90 nucleotides using FastqMcf (70). Further quality control steps involved the removal human reads and the reads from the Illumina spike-in (bacteriophage phiX174), by mapping the reads against the corresponding genomes with Bowtie2 (71). Metatranscriptomes were additionally processed to remove ribosomal RNA, by mapping the reads against 16S and 23S rRNA gene databases (SILVA_119.1_SSURef_Nr99_tax_silva and SILVA_119_LSURef_tax_silva (72)), and to remove contaminant adapters using trim_galore (http://www.bioinformatics.babraham.ac.uk/projects/trim_galore/) with parameters: -q 0, --nextera, and --stringency 5. The milk sample of mother-infant Pair 4 at Timepoint 1 was discarded from further analyses because of the low number of microbial reads (less than 400,000 bp) obtained after the quality control (**Supplementary Table S1**). All metagenomes and metatranscriptomes have been deposited and are available at the NCBI Sequence Read Archive under the BioProject accession number PRJNA339914.

### Taxonomic and strain-level analysis

Taxonomic profiling was performed with MetaPhlAn2 (45) (with default parameters) on all the 24 metagenomic samples sequenced. MetaPhlAn2 uses clade-specific markers for taxonomically profiling shotgun metagenomic data and quantify the clades present in the microbiome with specie-level resolution.

Strain-level profiling was performed with PanPhlAn (29) and a novel strain-level method called StrainPhlAn. PanPhlAn is a pangenome-based approach that profiles the presence/absence pattern of species-specific genes in the metagenomes. The genes presence/absence profiles are then used to characterize the strain-specific gene repertoire of the members of the microbiome. PanPhlAn has been executed using the following parameters: --min_coverage 1, --left_max 1.70, and --right_min 0.30. StrainPhlAn is a novel complementary method based on SNVs analysis that reconstructs the genomic sequence of species-specific markers. StrainPhlAn builds the strain-level phylogeny of microbial species by reconstructing the consensus marker sequences of the dominant strain for each detected species. The extracted consensus sequences are multiple-aligned using MUSCLE version v3.8.1551 (73) (default parameters) and the phylogeny is reconstructed using RAxML version 8.1.15 (74) (parameters, -m GTRCAT and -p 1234). StrainPhlAn is available with supporting documentation at http://segatalab.cibio.unitn.it/tools/strainphlan.

### Functional profiling from metagenomes and metatranscriptomes

The functional potential and transcriptomic analysis was performed with both HUMAnN2 (75) and PanPhlAn (29).HUMAnN2 selects the most representative species from a metagenome and then builds a custom database of pathways and genes that is used as a mapping reference for the coupled metatranscriptomic sample to quantify transcript abundances. We computed the normalized transcript abundance (NTA), that we define as the average coverage of a genomic region in the metatranscriptomic versus the corresponding metagenomic sample normalized by the total number of reads in each sample. PanPhlAn infers the expression of the strain-specific gene families by extracting them from the metagenome and matching them in the metatranscriptome. PanPhlAn has been executed using the following parameters: --rna_norm_percentile 90 and --rna_max_zeros 90.

### Profiling of DNA and RNA viruses

We investigated the presence of viral and phage genomes by mapping the reads present in the metagenomes and metatranscriptomes against 7,194 viral genomes available in RefSeq (release 77). The average coverage and sequencing depth were computed with samtools (76) and bedtools (77).

The presence of the Pepper Mild Mottle Virus (PMMoV) was confirmed by mapping the reference genome (NC_003630) against the metatranscriptomic samples from mother-infant of Pair 4 and Pair 5. In the mother and infant of Pair 4, 424,510 and 119 reads were mapped respectively, while in the mother and infant of Pair 5, 1,444 and 61 of the reads were mapped respectively. In the two mothers (Pair 4 and 5) the breadth of coverage is 0.99 and 0.98, and the average coverage is 6,562 and 22, respectively. In the two infants (Pairs 4 and 5), the breadth of coverage is of 0.6 and 0.5, and the average coverage is of 1.81 and 0.95, respectively. Additionally, we extracted the shared fractions of the PMMoV genome present in both mother and infant of Pair 4, together with the same regions of all the available reference genomes (n=13 and specifically accessions LC082100.1, KJ631123.1, AB550911.1, AY859497.1, KU312319.1, KP345899.1, NC_003630.1, M81413.1, KR108207.1, KR108206.1, AB276030.1, AB254821.1, LC082099.1).The resulting sequences were aligned using MUSCLE version v3.8.1551 (default parameters) and the resulting alignment was used to build a phylogenetic tree with RAxML v.8.1.15 (parameters, -m GTRCAT and -p 1234).

### Statistical analyses and data visualization

The taxonomic and functional heatmaps were generated using hclust2 (parameters: --f_dist_f euclidean, --s_dist_f braycurtis, and -l) available at https://bitbucket.org/nsegata/hclust2. The Multi-Dimensional Scaling plots were computed with the sklearn Python package (78). Biomarker discovery (**Supplementary Fig. S4)** was performed by applying LEfSe (79) (with parameter: -l 3.0) on HUMAnN2 profiles. The two functional trees (**Supplementary Fig. S4**) have been automatically annotated with export2graphlan.py (GraPhlAn package) and displayed with GraPhlAn (80) using default parameters.

## Acknowledgements

We would like to thank prof. Marco Ventura and his group for performing the DNA extraction from the milk samples. This work was supported by Fondazione CARITRO fellowship Rif.Int.2013.0239 to N.S. The work was also partially supported by the People Programme (Marie Curie Actions) of the European Union Seventh Framework Programme (FP7/2007-2013) under REA grant agreement no. PCIG13-GA-2013-618833 (N.S.), by startup funds from the Centre for Integrative Biology, University of Trento (N.S.), by MIUR “Futuro in Ricerca” RBFR13EWWI_001 (N.S.), by Leo Pharma Foundation (N.S.), and by Fondazione CARITRO fellowship Rif.int.2014.0325 (A.T.).

